# Generation of a novel growth-enhanced and reduced environmental impact transgenic pig strain

**DOI:** 10.1101/250902

**Authors:** Xianwei Zhang, Zicong Li, Huaqiang Yang, Dewu Liu, Gengyuan Cai, Guoling Li, Jianxin Mo, Dehua Wang, Cuili Zhong, Haoqiang Wang, Yue Sun, Junsong Shi, Enqin Zheng, Fanming Meng, Mao Zhang, Xiaoyan He, Rong Zhou, Jian Zhang, Miaorong Huang, Ran Zhang, Ning Li, Fanming Zhe, Jinzeng Yang, Zhenfang Wu

**Affiliations:** College of Animal Science, South China Agricultural University, Guangzhou, China.; National Engineering Research Center for Breeding Swine Industry, Guangdong Wens Foodstuff Group Co., Ltd, Yunfu, China.; Department of Animal Biosciences, University of Guelph, Guelph, Ontario, Canada.; Department of Human Nutrition, Food and Animal Sciences, University of Hawaii at Manoa, Honolulu, HI, USA.; College of Biological Science, China Agricultural University, Beijing, China.

## Abstract

In pig production, insufficient feed digestion causes excessive nutrients such as phosphorus and nitrogen, which are then released to the environment. To address the issue of environmental emissions, we have established transgenic pigs harboring a single-copy quad-cistronic transgene and simultaneously expressing three microbial enzymes, β-glucanase, xylanase, and phytase in the salivary glands. All the transgenic enzymes were successfully expressed, and the digestion of non-starch polysaccharides (NSPs) and phytate in the feedstuff was enhanced. Fecal nitrogen and phosphate outputs were reduced by 23%–46%, and growth rate improved by 23.4% (gilts) and 24.4% (boars) when the pigs were fed on a corn and soybean-based diet and high-NSP diet. The transgenic pigs showed a 11.5%– 14.5% improvement in feed conversion rate compared to the age-matched wild-type littermates. These findings indicate that transgenic pigs are promising resources for improving feed efficiency and reducing nutrient emissions to the environment.

## Introduction

Annual global pig production is approximately 1.2 billion heads, with more than half produced in China (***USDA, 2016***). Grains are the main feedstuff of the pig industry; however, its production capacity in China and many other countries is insufficient. For example, China imported 254.08 million metric tons of soybean and 59.28 million metric tons of coarse grain between 2014 and 2016 for pig production. This accounts for 63.3% and 11.2% of the annual global Imports volume, respectively (***USDA, 2017***). The livestock industry often allows animals to achieve maximal growth in order to fully utilize their economic outputs, yet ineffective feed digestion can cause serious nutrient emissions to the environment. Two nutrients that have received the most attention from environmental groups are nitrogen (N) and phosphorus (P) and are often supplied in excessive amounts in the diet for maximal growth. In pig production, only 1/3 of feed N and P is metabolically utilized in cereal and soybean-based diets. The deposition rate of N is only 25%–32% in grower-finisher pigs (***Shirali et al., 2012)***. It has been reported that N excretion is up to 20 kg/sow/year and 25 kg/boar/year (***DEFRA, 2007***). Approximately 51% of N intake is excreted in urine, which is mainly from protein metabolism and underutilized amino acids and non-protein nitrogen (NPN) (***Shirali et al., 2012)***. Fecal N excretion comes from undigested protein fractions and endogenous tissue losses such as digestive enzyme secretions and desquamation of intestinal cells, which accounts for 17% of the N intake. Only approximately 30% of P is retained in a grower-finisher pig on a cereal-soybean meal-based diet. In total, 70% of ingested P is excreted either through the feces or urine (***Dourmad et al., 1999)***. The total P emission of a grower pig (weight range: 0–108 kg) is 1.35 kg per year, accounting for 67.2% of the total P intake. A productive sow emits 5.42 kg P per year, accounting for 77.7% of the total P intake (***Dourmad et al., 1999***). The N and P from animal excreta are the main pollutants of the water, soil, and air of intensive pig production sites. Surface water becomes eutrophic when excessive P and N inputs, thereby causing overgrowth of blue-green algae and death of aquatic animals (***Jongbloed and Lenis, 1998***; ***Poulsen***, ***1998***). Considering all these aspects, improving nutrient utilization in feed is of great significance to maximize feed grain utilization as well as for environmental conservation.

Non-starch polysaccharides (NSPs) consist of a series of soluble and insoluble polysaccharides that are primarily present in plant cell walls (***McDougall, 1996***; ***Sarkar et al., 2009***). In cereal grains, arabinoxylans and β-glucans are found in the cell walls of the protein-rich aleurone layer and starchy endosperm and can act as barrier against nutrient hydrolysis and absorption (***Stone***, ***1981***). Similarly, the cell wall polysaccharides of soybean, canola seed, and peas may also be responsible for this nutrient-encapsulating effect (***Omogbenigun et al., 2004***). Therefore, NSPs are the main anti-nutrient factors of cereal and bran (***Fangel et al., 2012***; ***Sarkar et al., 2009***). Pigs inherently lack NSP-degrading enzymes (NSPases) in their digestive tracts. Pigs are inherently incapable of digesting NSPs (***Hooda et al., 2010***), but can be partly degraded by the natural microbial community thriving in the intestinal tract. The unused NSPs restrict the utilization of organic material, crude protein, and energy of feed grain (***Högberg and Lindberg, 2004)***. High-P emission from monogastric animals such as pigs and poultry arises from their poor physiological ability to hydrolyze plant phytates, which account for up to 80% of P in common cereal grains, oil seed meals, and by-products (***Sivaloganb, 1994***). Phytates are negatively charged saturated cyclic acids that can bind to positively charged molecules in the diet such as minerals and protein, thereby reducing nutrient digestibility and increasing discharge of the unabsorbed nutrients to the environment (***Dersjant-Li et al., 2015***).

Various methods have been employed to address the issues of insufficient utilization of feed nutrients in the pig industry. For example, dietary supplementation of phytate- or NSP-degrading enzymes has been proposed to reduce P or N emissions from pig farms (***Golovan et al., 2001***; ***Barrera et al., 2004***; ***Golovan, et al., 2001***; ***Jongbloed and Lenis, 1998)***, as well as increasing pig body weight gain and feed conversion efficiency (***Diebold et al., 2004***; ***Willamil et al., 2012***; ***Woyengo and Nyachoti, 2011***). However, the effects of these enzyme supplements are distinct the phytates or NSP content of the diets and the type of enzyme. Adjusting the nutrient content of the diet such as lowering crude protein or supplementation of limiting amino acids can reduce N emissions from unutilized nutrients in feces and slurry (***Aarnink and Verstegen, 2007)***. Recent advancements in genetic engineering and animal cloning technologies have facilitated in the establishment of genetically modified pigs with economically significant traits. Transgenic (TG) pig lines secreting salivary bacterial phytases have been generated (***Kiarie et al., 2012)***. The P content of fecal matter from TG weaner and grower-finisher pigs fed on soybean meals has decreased by as much as 75% and 56%, respectively, compared to their non-TG counterparts. Endogenous salivary phytase significantly promotes the digestion of P from dietary phytates (***Kiarie et al., 2012)***. To our knowledge, no pig lines that express NSP-degrading enzymes have been establishment to date. In this study, we established stable transgenic pig lines that co-expressed NSP-degrading enzymes (β-glucanase and xylanase) and phytase in saliva. We also show that these TG pigs are capable of transmitting transgenes to their progeny. Multiple enzymes coordinately degrade NSPs and phytates in feed grains. We also report the grain digestibility, nutrient emission, growth performances, and feed conversion rate of the TG pigs compared to their wild-type controls.

## Results

### Optimization and construction of a 2A-mediated salivary gland-specific multi-transgene

Through characterization of multiple codon-optimized β-glucanase genes fused with the N-terminal porcine parotid secretory protein (PSP) signal peptide, we have determined that BG17A and EG1314 exhibited optimal activity and stability in porcine cells, and the pH condition is compatible to that of the pig digestive tract (***Zhang et al., 2015***). We previously reported that the fused BG17A and EG1314, which was linked by a self-cleaving 2A peptide, had a broader optimal pH range and higher stability in an acidic environment than either of these alone (***Zhang et al., 2015***). After codon optimization and fusion with the pig PSP signal peptide, three xylanases (XYNB, XYL11, and XYF63 (also known as XYN11F63) were transfected into PK15 cells and subjected to enzymatic activity assay. Among the three xylanases, XYNB presented the highest enzymatic activity (***Supplemental figure 1A***) and stability (***Supplemental figure 1B***). In addition, XYNB showed greater resistance to peptic and tryptic hydrolysis than the other two xylanases (***Supplemental figures 1C-E***). As for the two phytases, *Citrobacter freundii* APPA (CAPPA) only had two narrow peaks at the optimal pH levels of 2.5 and 5.0, respectively (***Supplemental figure 2A***), whereas *Escherichia coli* APPA (EAPPA) exhibited a broad optimal pH ranging from 1.5 to 5 (***Supplemental figure 2B***). EAPPA was more tolerant of pepsin and trypsin than CAPPA. There was almost no reduction in activity of EAPPA after a 2-h pepsin treatment, whereas 52.2% of the biological activity was left after CAPPA treatment (***Supplemental figure 2C***). When treated with trypsin alone, EAPPA and CAPPA retained 98.2% and 39.7% of its activity, respectively; when treated with trypsin + EDTA, EAPPA and CAPPA retained 31.8% and 13.7% of its activity, respectively (***Supplemental figure 2D***). Based on these results, two β-glucanases genes (*bg17*A and *eg1314*), a xylanase gene (*xyn*B), and a phytase gene (e*appA*) showed better performance that the other candidate transgenes.

A polycistronic cassette of fusion enzymes was constructed by head-to-tail ligation of four selected genes and flanked on the 3’ end by an Hemagglutinin (HA) tag. The DNA sequences of the self-cleaving peptides E2A, T2A, and P2A were used as linkers between the coding DNA sequences of two neighboring enzymes (***Supplemental figure 3A***). Before construction of the final TG vector, the fusion enzyme sequences were ligated downstream of the CMV promoter, and the expression level and enzymatic activity of each fusion enzyme were measured in porcine cells. We were able to detect the expression of all the four active enzymes in the PK15 cells, although the expression level and enzymatic activity of each recombinant fusion enzyme was lower than its original monomeric counterpart (***Supplemental figures 3B and C***). The lower transfection efficiency of the large transgene construct likely accounted for the observed lower expression and enzymatic activity in cells. We then employed a mouse PSP promoter to replace the CMV promoter to control the salivary gland-specific expression of the fusion gene, and this expression cassette was inserted, together with a CMV promoter-driven neo-EGFP fusion gene, into the piggyBac transposon vector to form a transgenic vector, namely, pPB-mPSP-*BgEgXyAp*-*neoGFP* (***Figure 1A***).

**Figure 1.**
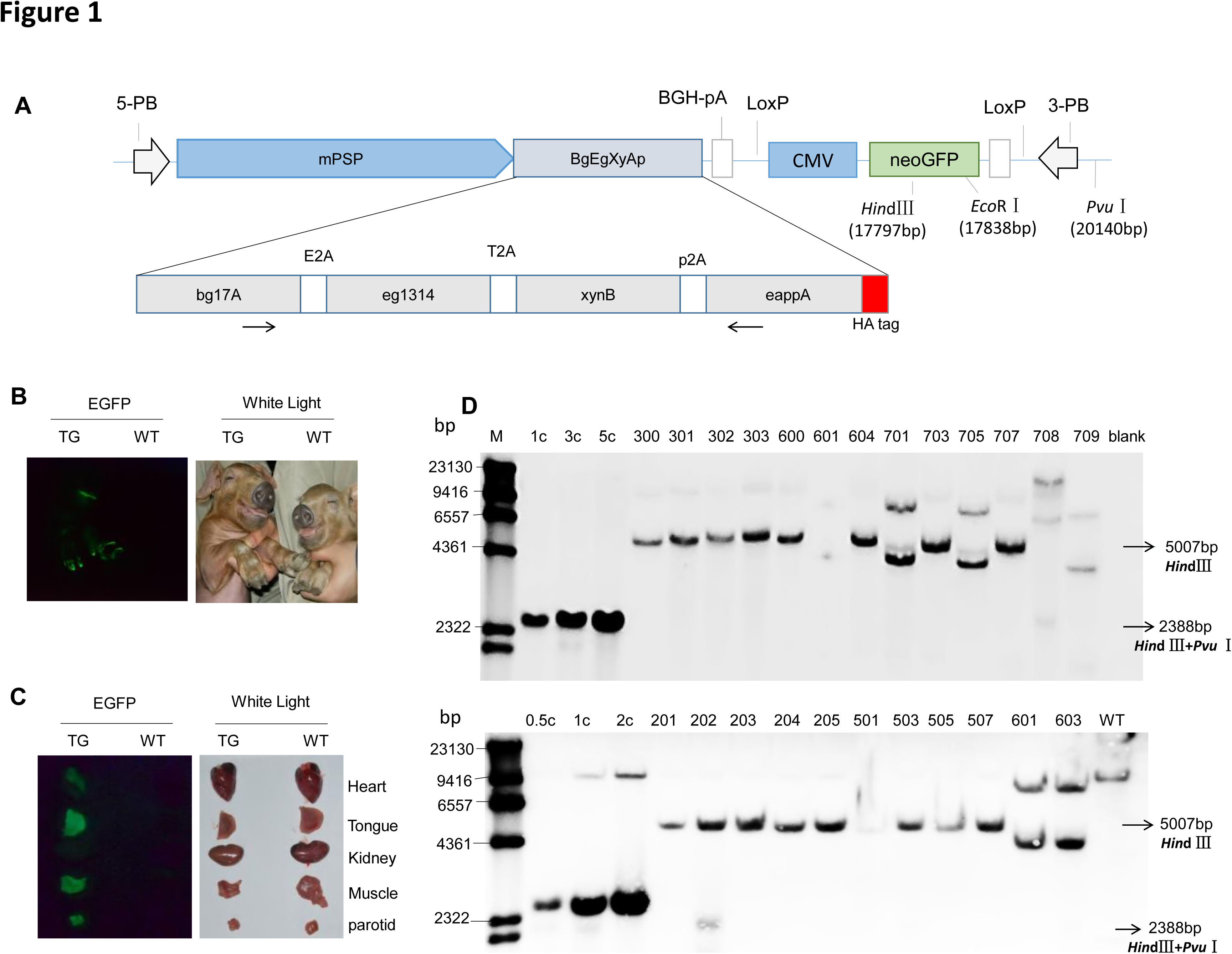
(***A***) DNA construct that was integrated into the piggenome for expression of the transgenic fusion enzyme in saliva. mPSP: Mouse parotid secretory protein promoter. BGH: Bovine growth hormone polyadenylation signal. Total length: 19,886 bp. (***B***) Expression of EGFP in the whole body of TG pigs. (***C***) Expression of EGFP in the heart, tongue, kidney, muscle, submandibular gland, spleen, lung, and liver of TG pigs. (***D***) Southern blot analysis of multi-enzyme transgene integration in TG pigs. 0.5c, 1c, 2c, 3c, and 5c represent copy number of transgenic vector used as loading controls. The probe is shown in Figure1A. Blank: Blank control (ddH_2_O).

### Generation of TG pigs

The resulting TG piggyBac transposon vector (pPB-mPSP-*BgEgXyAp*-*neoGFP*) and a piggyBac transposase expression vector (hyPBase) were co-transfected into the porcine fetal fibroblasts (PFFs) of a male Duroc pig. Transfected PFFs were selected with G418 for approximately two weeks, and the resulting EGFP-expressing cell colonies were pooled and identified by PCR for the presence of the transgene. The positive cell colonies were used as donor cells for somatic cell nuclear transfer. A total of 4,008 reconstructed embryos were generated and transferred to 16 recipient sows (***Supplemental table 13***). A total of 35 cloned male Duroc piglets were born, of which 24 founders were positive for transgene by PCR detection (***Supplemental figures 4A-B***-***figure supplement 1-2***). Bright green fluorescence signals were observed in the entire body and various organs of the piglets (***Figure 1B-C-figure supplement 1***). Eight of the transgenic pigs survived to sexual maturity. The *hyPBase* gene was not detected in the genome of the TG pigs. Among the 24 TG founders, four piglets (601, 701, 705, and 709) harbored the fragments of the ampicillin-resistance gene of the transgene vector (***Supplemental figures 4A-figure supplement 1***), implying the occurrence of unexpected breaks within the transgene. The other 20 piglets harbored the intact transgene expression cassette (total length: 19,886 bp). Southern blotting, quantitative PCR, inverse PCR, and sequencing results further demonstrated that 19 piglets carried a single copy of the transgene (***Figure 1D-figure supplement 2, Supplemental figures 4C-figure Supplement 2, and Supplement figure 5d***), of which two carried a single copy of the transgene that was inserted into intron 1 of *Legumain* (line 1) (***Supplement figure 4C-figure supplement 2, Supplement table 1***); 17 carried a single copy of the transgene that was integrated into intron 5 of *CEP112* (line 2, of which six piglets survived) (***Figure 1D- figure supplement 2***;***Supplement table 1*)**. One (708) carried three copies, and the integration site was the intergenic region between *LOC100525528* and *CXCL2* (***Figure 1D-figures supplement 2*** *and* ***Supplemental figures 5***;***Supplemental table 1***).

RT-PCR analysis indicated that the BgEgXyAp, bg17, eg1314, xynB, and eappA transgenes were unambiguously expressed in the parotid, submandibular, and sublingual glands, whereas these were undetectable in the other tissues of the TG founders such as the lungs, heart, liver, stomach, spleen, kidney, duodenum, colon, and muscle (Figure 2A; Supplemental figure 1, 2, and 6). Quantitative PCR analysis indicated that the highest *BgEgXyAp* transgene expression levels were observed in the parotid gland, followed by the submandibular and sublingual glands, and trace or undetectable level were observed in the other tissues of the TG founders (***Supplemental figure 6***). Ectopic expression was not observed in this study. Western blot analysis demonstrated the expression of β-glucanase, xylanase, and phytase in the saliva of the TG founders (***Figure 2B-figure supplement2***). During the feeding period, the TG pigs produced 0.3-2.3 U/mg of β-glucanase, 0.6-2.4 U/mg of xylanase, and 0.5-5.7 U/mg of phytase in the saliva (***Figures 2C-E***). The total salivary protein concentrations of the TG and WT pigs are shown in ***Figure 2F***.

**Figure 2.**
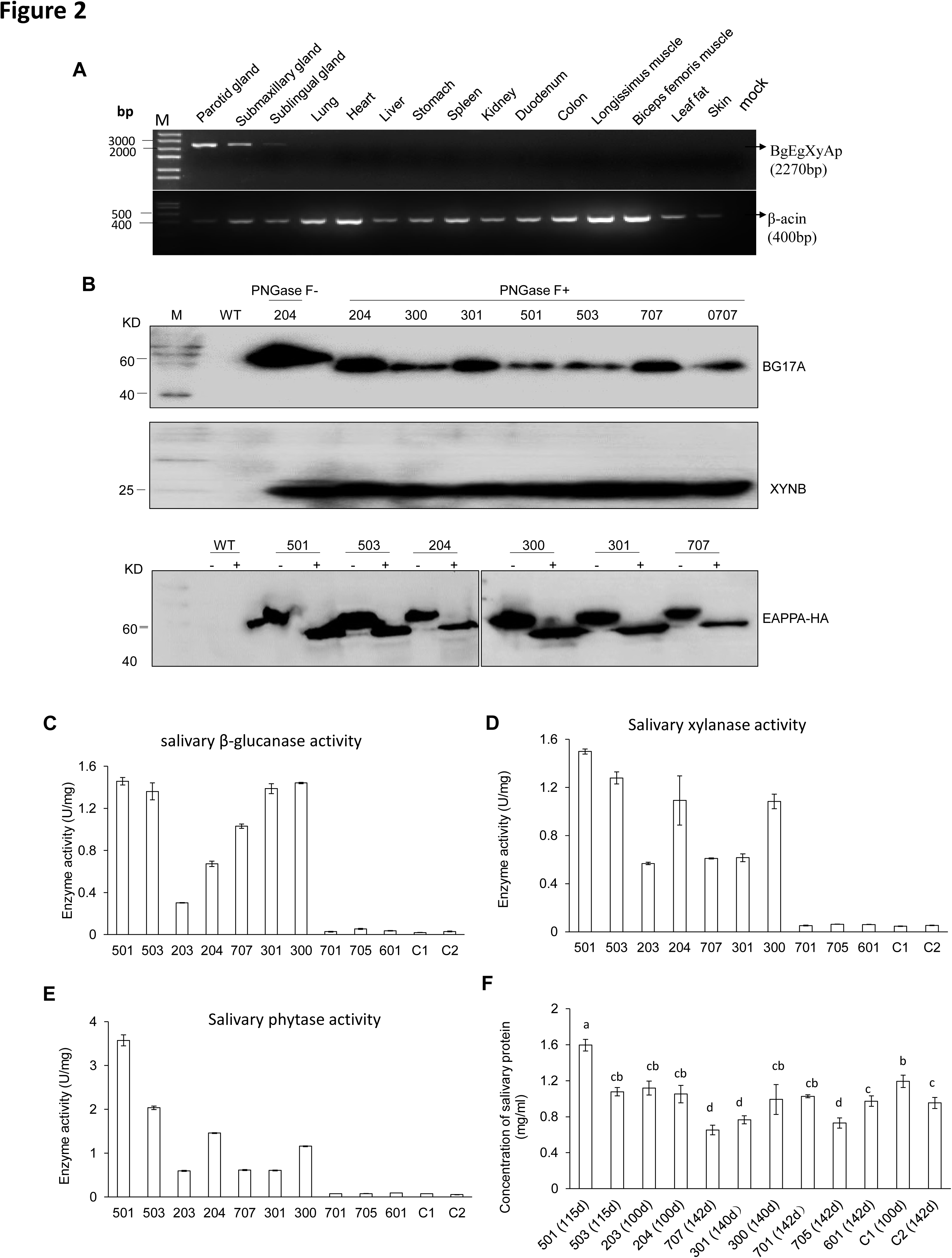
(***A***) RT-PCR assay for mRNAexpression profiles of transgenes in different tissues. Forward primers and reverse primer are bound to the *bg17A* gene and *eappA* gene, respectively (arrows are shown in Figure1A). Mock: Blank control (ddH2O). (***B***) Western blotting assay demonstrating the expression of BG17A, XYNB, and EAPPA in the saliva of TG pigs. Saliva samples were either incubated with PNGase F (+) or mock (-)-treated prior to western blotting to analyze the glycosylation status of the transgenic enzymes. PNGase F: Peptide N-glycosidase F. (***C-E***) Salivary β-glucanase, xylanase, and phytase activity assays of the TG pigs. (***F***) Concentration of total salivary protein of the TG and WT pigs. C1,C2: Age- and body weight-matched WT pigs. Data are expressed as the mean ± SEM, n = 3, a, b, c, and d values on the bar graph with different superscript letters differ significantly (one-way ANOVA, P < 0.05).

Pigs of TG line 2 that harbored a single-copy transgene within *CEP112* intron were used in growth trials and feed evaluations. The TG line 2 pigs were crossed with WT Duroc pigs, which generated 116 F1 progeny, of which 57 tested positive for the transgene. Four hundred and four of the F2 progeny were sired, of which 231 were positive for the transgene.

### Measurement of enzyme production in TG pigs

To understand the effect of the three enzymes of the TG pigs on nutrient digestion, we investigated the pattern of salivary secretion and enzyme production in the TG pigs. Saliva was collected from the unilateral parotid glands of the TG pigs and analyzed in terms of enzyme yield. The average β-glucanase, xylanase, and phytase yields were 2,331.84, 2,413.38, and 2,935.19 U per kilogram meal, respectively, in grower pigs, and 920.82, 939.03, and 1,042.19 U per kilogram meal, respectively, in finisher pigs (***Supplementaltable 2***). The volume of saliva collected from the parotid gland of the finisher pigs was significantly lower compared to that of the grower pigs, which may be attributable to the shorter feeding time (Ft) in finisher pigs. Furthermore, saliva and enzyme production at different time points were evaluated. The results showed that the pigs only secreted saliva from parotid gland during Ft. At the other time points, including 10 min before or after feeding (Bf or Af), insignificant amount of saliva was secreted (***Supplemental figure 7A***). The enzymes expressed by the transgene showed high enzymatic activity at Bf, Ft, and Af. At rest time (Rt), the saliva collected from either parotid or mouth showed reduced enzymatic activities. Of note, a significantly lower the enzyme activity was observed in the saliva samples collected at Rt (***Supplemental figure 7B***).

### Improved feed utilization and reduced nutrient emission in TG founders

The grower-finisher founder TG pigs (weight range: 30 kg to 50 kg) were fed corn-soybean (CS) or wheat-corn-soybean-bran (WCSB) diets (***Supplemental table 3***) to investigate the effects of a salivary cocktail of β-glucanase, xylanase, and phytase on feed utilization. The traditional CS diet contains a low level of NSPs and total P with a high proportion of phytates (70%), and the WCSB diet contains a relatively high concentration of NSPs and total P with 63% phytates. For each diet, six TG pigs and six age-matched and weight-matched non-TG Duroc boars (WTs) were fed; and six WTs were fed the same diet with supplementary multi-enzyme preparations of β-glucanase, xylanase, and phytase. After dietary treatment, nutrient digestion among the experimental groups was measured and compared. For both diets, the apparent total tract digestibility (ATTD) of dry matter (DM), P, N, and calcium (Ca) significantly increased in TG pigs compared to that of WT pigs (*Figure 3A*). The fecal outputs of N and P, relative to input by feed, significantly decreased in the TG pigs compared to that of the WT pigs (*Figure 3B*). Fecal N and P excretion decreased by 25% and 46%, respectively, with the CS diet, and 23% and 35%, respectively, with the WCSB diet. A significant reduction in total P and Ca (feces plus urine) was also observed in the TG pigs with both diets. Almost all tested items of the TG pigs showed some improvement compared to that of the WT(+) pigs that were fed the same diets supplemented with multi-enzyme preparations, although the differences were not statistically significant among groups (***Supplemental table 4 and 5***).

**Figure 3.**
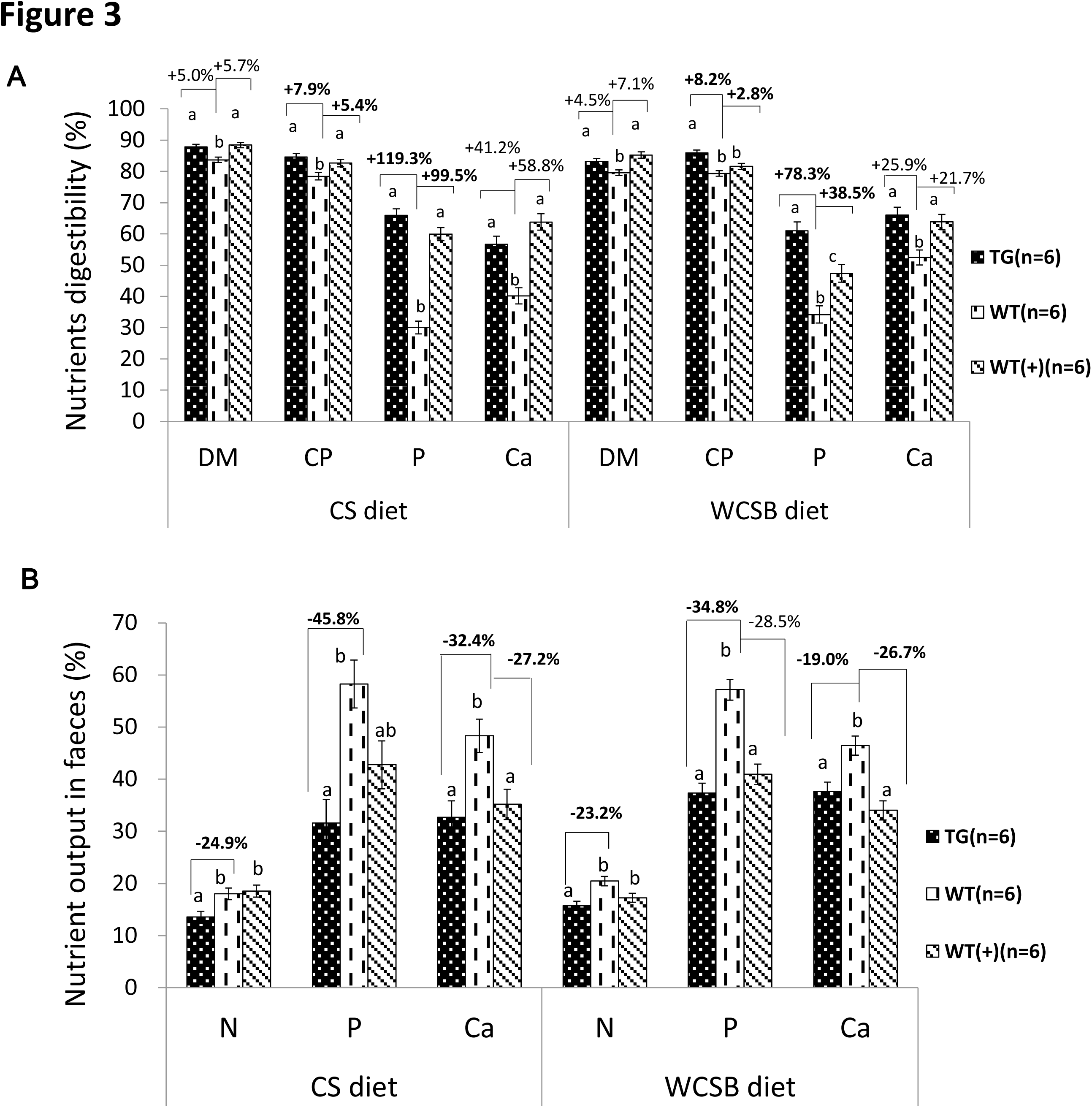
Comparison of the apparent total tract nutrient digestibility values (%) and fecal nutrient output (% of their dietary intake) between transgenic (TG) grower pigs and their wild-type (WT) littermates fed on corn and soybean meal (CS) and wheat- and corn and soybean meal(WCSB)-based diets with and without exogenous feed enzymes. (*A*) Comparison of the apparent total tract nutrient digestibility values (%) of dry matter (DM), crude protein (CP), phosphorus (P), and calcium (ca). (*B*) Comparison of fecal N, P, and Ca output. WT(+): WT grower pigs fed on the CS and WCSB diets supplemented with an optimal dose of β-glucanase, xylanase, and phytase. Data are expressed as the least square means (Lsmean ± SEM). ^a,b,c^ Values on the bar graph with different superscript letters differ significantly (ANCOVA, P < 0.05).

The serum components of the TG pigs fed on an LNHP diet containing a low N level and high proportion of phytates (78%) were analyzed (***Supplemental table 6***). Serum alkaline phosphatase activity in the TG pigs was lower than the WT littermates. Serum P and glucose levels of the TG pigs were greater than that of the WT littermates. The serum D-xylose levels of the TG pigs were slightly higher than the WT littermates. No differences in serum Ca, Zn, urea N, uric acid, and total protein concentrations between TG and WT pigs were observed.

### Enhanced growth performance in TG pigs

The growth performance of eight F1 TG pigs (females) and 17 WT littermates (females) were fed a low-N level and high proportion of phytates (LNHP) diets (***Supplemental table 7***) during the growing period from 30 kg to 50 kg weights was measured. The TG pigs exhibited a higher average daily gain (ADG) rate and lower feed conversion rate (FCR) than the WT pigs during this stage (***Supplemental table 8***). Growth performance of the F2 TG pigs was also measured. A total of 74 F2 TG pigs (23 boars, 51 gilts) and 52 WT littermates (21 boars, 31 gilts) were raised together and fed the same diets shown in ***Table supplement 9***. In this study, TG boars showed increased ADFI (P =0.077) compared to the WT boars that were fed the same diets (***Supplemental table 7A***). Similar results were observed in the gilts. Significantly improved ADG rates and lower FCR were observed in the TG boars and gilts compared to the WT gilts during the entire feeding period **(*Figures 4B and 4C*)**. It took an average of 110 days for TG boars to grow from 30 kg to 115 kg, whereas the WT boars needed 145 days of feeding on the same diets. Similar results were observed in the gilts. It took 121 days for the TG gilts to grow from 30 kg to 115 kg, whereas the WT females required 150 days (*Figure 4D*). Taken together, the TG pigs showed a 7.0%-7.6% higher ADFI than the WT pigs over the same grower and finisher phases (30 kg to115 kg), but gained 23.0%-24.4% more body weight (BW) daily than the WT pigs over the same grower and finisher phases (30 kg to 115 kg). The time to reach 115 kg BW was shortened by 19.2%-21.9% (29.6 days to 35.1 days). The FCR decreased by 11.5% -14.5% during the grower and finisher periods.

**Figure 4.**
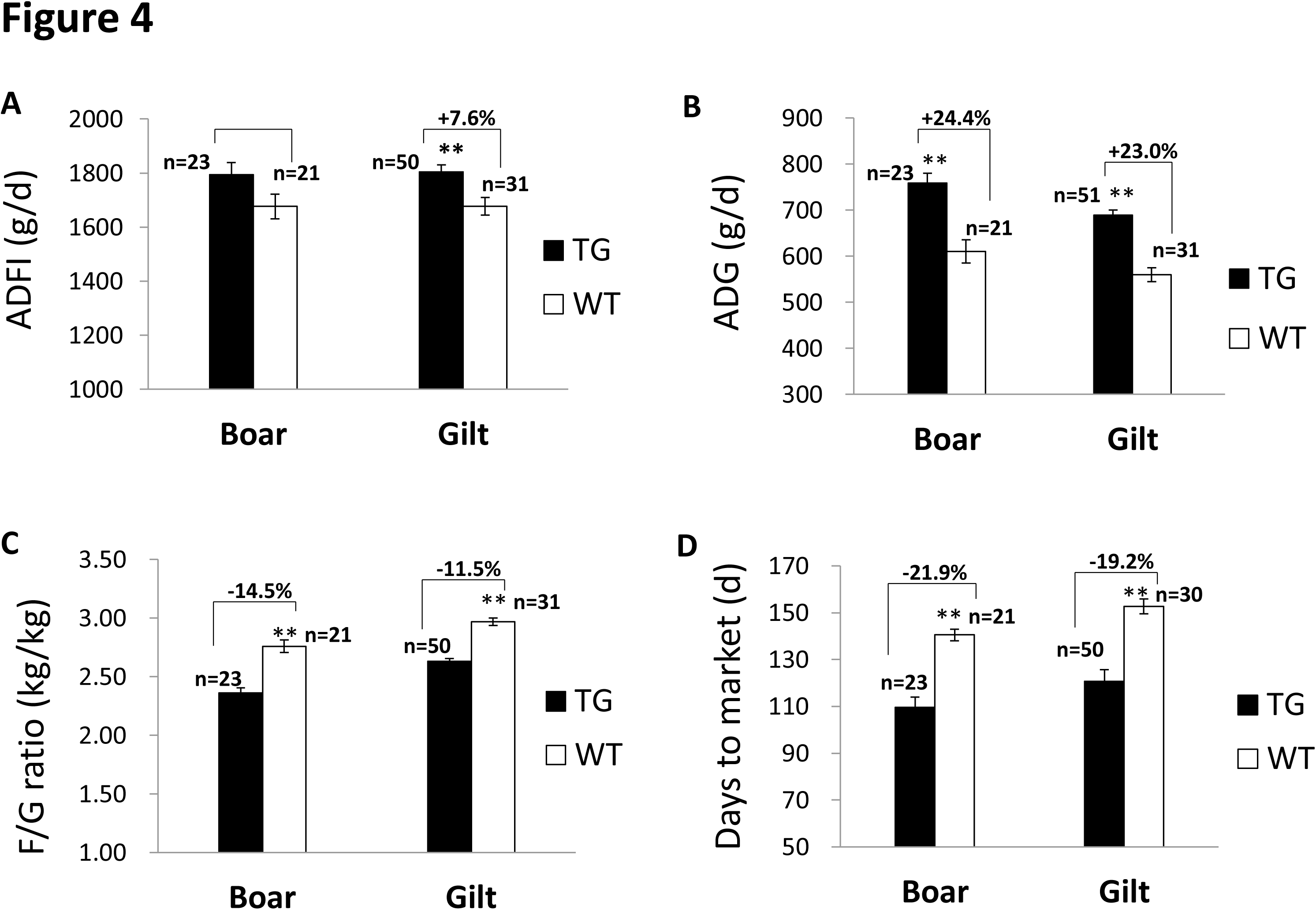
(***A***) Comparison of average daily feed intake (ADFI). (***B***) Comparison of averagedaily gain (ADG). (***C***) Comparison of feed/gain (F/G). (***D***) Comparison of days to market. Data are expressed as the least square means (Lsmean ± SEM), asterisks indicate significant differences between TG and WT pigs within one line (ANCOVA, **P < 0.01).

## Discussion

Previous studies have shown that the salivary gland secretes a single microbial enzyme phytase, and that pig manure has reduced P levels (***Kiarie et al., 2012***; ***Meidinger et al., 2013)***. The major objective of this study was to enhance the digestive utilization of feed grain and decrease the N and P emissions from pig manure. By employing an animal cloning procedure, we incorporated three microbial enzyme genes, namely, glucanase, xylanase, and phytase, into the pig genome. This is the first report that describes the integration of multiple single-copy genes into the pig genome. We obtained TG pigs by introducing the microbial genes *bg17A*, *eg1314*, *xynB*, and *eappA*, which encode for endo-β-1,3-1,4-glucanase, endo-xylanase, and 6-phytase, respectively, into the genome of pigs so that these could produce phytate- and NSP-degrading enzymes. These enzymes, which are inherently secreted by microbial communities, were optimized to adapt to the digestive tract environment of pigs and expressed specifically in the porcine salivary gland to initiate the digestion process of NSPs and phytates in the mouth. The feeding trials demonstrated that the TG pigs possessed a high digestive capacity for N, DM, phytate P, and other nutrients, enhanced growth performance, and decreased manure nutrient emissions. We did not find any negative side effects in these TG animals such as changes in spirit, behavior, reproductive capacity, viability, growth performance, blood physiology, and biochemistry.

The viral 2A peptide, which is called a ‘‘self-cleaving’’ peptide or protease site because it can be cleaved at the C terminus between the last two amino acids through ribosomal skipping during protein translation, results in the expression of two independent proteins from a co-transcribed fused mRNA. Previous studies have shown that F2A can successfully mediate bicistronic expression, but the co-expression of the four genes mediated by F2A was inefficient in the cultured cells and embryos. The expression of the last two genes was much weaker than that of the first two genes in the polycistronic cassette (***Deng et al., 2011***; ***Kim et al., 2011***). There are several types of 2A peptides in various species. Among these, P2A has the highest cleavage efficiency in zebrafish, followed by T2A, E2A, and F2A (***Kim et al., 2011***). The present study constructed quad-cistronic genes, which were mediated by three types of 2A (E2A, P2A, and T2A), and driven by a PSP promoter. In previous studies, the cleavage efficiency of a 2A peptide was estimated to be within the range of 80% to 95%. In the present study, the three 2A peptides almost completely trimmed the four transgenes. The ‘‘cleavage’’ at a 2A site results in an upstream protein that is fused to the 2A peptide without a proline residue and a downstream protein with a proline residue added tothe N terminus, which has the potential to significantly affect its function (***Deng et al., 2011***; ***Tang et al., 2009***). No adverse effects on the function of target protein due to the fusion of the 2A residue were observed in this study.

In this study, transgenes encoding microbial enzymes were driven by the salivary gland-specific PSP promoter. TG pig line 2 harbors a single copy of a multi-transgene, which facilitates the establishment of a breeding population with identical genotypes. Compared to the previously reported phytase TG pig lines that harbor a high copy number of a foreign gene in an insertion locus (2 to 35 copies of the transgene, producing 5 to 6,000 U/mL of phytase) (***Golovan et al., 2001)***, our multi-TG pigs secreted a lower amount of enzymes into the saliva. In fact, our multi-TG pigs demonstrated similar digestibility with the high-expression phytase TG pigs, as well as reduced emission of P in manure. For grower-finisher pigs, our study displayed an approximately 45.7% reduction in fecal P when fed CS meals (70.3% phytates), versus 56% reportedly in high-expression phytase TG pigs fed on soybean meals (53% phytates) (***Forsberg et al., 2013***; ***Golovan et al., 2001***). The three major salivary glands showed high expression levels and enzymes activities, which could be arranged in decreasing order as follows: parotid gland > sublingual gland > submaxillary glands, which agree with the findings of a previous report (***Golovan et al., 2001)***. The enzyme activities per milliliter of parotid saliva between grower pigs and finisher pigs were similar. The expression levels of glucanase, xylanase, and phytase in the bilateral parotid glands of the TG pigs were 4,663.68, 4,826.76, and 5,870.38 U/kg diet intake for grower pigs, and 1,841.64, 1,878.06, and 2,084.38 U/kg diet intake for finisher pigs. These levels of expression are higher than the amounts used as dietary supplements.

The TG pigs demonstrated increased P retention and reduced percentages of manure P excretions compared to age-matched WT pigs fed on the same diets, implying the significant digestive effect of TG phytase. The P retention rate was 54.7%-67.3%, and reduction in total P output ranged from 21.29% to 44.21%. These results are slightly lower than that of previously reported TG phytase pigs, which had P retention rates of 57.2%-77.8%, and total P output decreased by 27.5%-62.0% (***Golovan et al., 2001***; ***Meidinger et al., 2013***). This discrepancy may be caused by the different expression levels of transgenes and the ingredients of the diets. However, the new pig model demonstrated that N digestibility was significantly enhanced in all diets tested. WCSB diets contain high levels of NSPs, which account for the highest fraction of polymeric carbohydrate in protein-rich materials (aleurone layer). Glucanase and xylanase secreted by TG pigs can effectively degrade the glucans and xylans in the cell walls of the aleurone layer, in which the matrix proteins and phytate globoids would be exposed and subjected to degradation by phytase and endogenous proteases. These results agree with that of high-NSP diets supplemented with NSPase (***Diebold et al., 2004***; ***Willamil et al., 2012***). For the corn-soybean diet, the effects of enzyme supplementation on nutrient digestibility of pigs, as reported in the literature, were highly variable (***Defa Li, 1999***; ***Ji et al., 2008***; ***Kim et al., 2003***; ***Willamil et al., 2012***). Some studies show positive responses to enzyme supplementation (***Ji et al.,2008***;***Kim et al.,2003***), whereas others do not (***Defa Li, 1999***; ***Willamil et al., 2012***). TG pigs fed on CS diets also showed significantly enhanced N digestibility. As the main ingredient of CS diets, corn contains high levels of phytates. The synergistic effect of NSPase (glucanase and xylanase) and phytase involves the digest of NSPs and phytates in the aleurone layer and endosperm cells of corn, thereby alleviating the anti-nutrition effect of phytates by reducing the binding of phytates to proteins and digestive enzymes in the digestive tract. In terms of N emission, the grower TG pigs emitted 24% less fecal N than the WT pigs. The NSPs in the cell wall are degraded by NSPases (glucanase and xylanase) that are secreted in the upper digestive tract. Moreover, NSPases can reduce the compensated secretion of endogenous fluids to decrease endogenous nutrient losses (***Kerr and Shurson, 2013***). TG phytase facilitates in the release of phytate-chelating proteases and other digestive enzymes to further degrade nutrients.

The TG pigs displayed better growth performance than conventional pigs fed on diets without supplemental P during the grower and finisher phases. Both TG boars and gilts showed a faster rate in BW gain, and exhibited greater feed efficiency than the respective WT littermates. Blood serum measurements also demonstrated enhanced serum P, glucose, and xylose levels in TG pigs, which signified enhanced digestive utilization of these nutrients. A significant decrease in serum alkaline phosphatase levels is an indicator of well-developed bone (***Meidinger et al., 2013***; ***Sefer et al., 2012***; ***Selle and Ravindran, 2008)***. These results are clear manifestations of the improved nutrient digestibility and growth performance of TG pigs.

In summary, the multi-TG pigs reported in the present study exhibited significantly enhanced digestibility of feed N, P, Ca, and other minerals. The pigs produced manure with significantly reduced P and N emissions into the environment as well as exhibited improved growth performances. This genetic strategy offers a very valuable biological solution for insufficient feed digestion and environmental emissions due to the global expansion of the livestock industry.

## Materials and methods

### Animals

Animal use followed the Instructive Notions with Respect to Caring for Laboratory Animals, issued by the Ministry of Science and Technology of China, and the NIH Guide for the Care and Use of Laboratory Animals. The animal use protocol was approved by the Institutional Animal Care and Use Committees (IACUCs) of South China Agricultural University.

### Transgene constructs

The *bgl7A*, *eg1314*, *xynB*, and e*appA* genes were fused in a head-to-tail tandem array, with E2A, T2A, and P2A used as linkers between them. Flag-tag and HA-tag were added to the C terminal of Eg1314 and EAPPA, respectively, to facilitate the detection of protein expression. The fusion genes were named *BgEgXyAp*, which was cloned into pCDNA3.1(+) to examine its expression and enzyme activity levels in porcine cells. A 12.1-kb upstream genomic sequence of murine PSP, as a promoter to drive the expression of the fusion gene specifically in salivary gland, were cloned and ligated to *BgEgXyAp*, and then introduced into the transposon piggyBac vector pPB-lox-*neoEGFP*-loxp (a gift from The Wellcome Trust Sanger Institute, Cambridgeshire, UK) to form the final transgene construct. The final construct was confirmed by sequence analysis.

### Transgenic pigs

Primary PFFs were isolated from 35-day-old male fetuses of Duroc pigs. PFFs were cultured in Dulbecco's modified Eagle's medium (DMEM, HyClone, Logan, UT, USA) supplemented with 12% fetal bovine serum (FBS, HyClone, Logan, UT, USA) and 1% (v:v) penicillin/streptomycin (10,000 U/mL penicillin, 10,000 μg/mL streptomycin; GIBCO-BRL, Grand Island, NY, USA) at 39°C in an incubator with 5% CO_2_. The transgene was mixed with a transposase, pCMV-hyPBase (a gift from the University of Hawaii, Honolulu, HI), and transfected into PFFs by electroporation (BTX, San Diego, CA). The transfected cells were split 1:6 into fresh culture medium. After 24 h, 300 μg/mL G418 (Gibco) was added to the medium to select transfected cell colonies, and the plates were incubated in media containing G418 for about 15 days. The surviving cell colonies with EGFP expression were selected and propagated in a fresh 24-well plate. Colonies that proliferated well were then expanded and screened for the presence of the *BgEgXyAp* transgene.SCNT was performed as previously described (***Deng et al., 2011***; ***Lai et al., 2006***). The reconstructed embryos were surgically transferred to the oviduct of the recipient gilts the day after estrus was observed. The pregnancy status of the surrogates was detected using an ultrasound scanner at 26 d after the embryo was transferred. This status was then monitored weekly before the expected due date. The cloned piglets were born by natural birth.

### Enzymatic activity assay

The supernatants from transfected cells and saliva from transgenic and non-transgenic pigs were used as total protein samples for the enzymatic activity assays. β-glucanase and xylanase activity assays were based on estimating the amount of reducing sugars released from the relevant substrates in the reactions using 3,5-dinitrosalicylic acid (DNS) reagent, as previously described (***Liu et al., 2010***; ***Luo et al., 2010***). One unit of activity was defined as the quantity of enzyme that releases reducing sugar at the rate of 1 μmol/min.

Phytase activity in saliva was determined by means of vanadium molybdenum yellow spectrophotometry. The reaction was performed in a final volume of 600-μL solution containing 0.25 M of acetate buffer (pH 5.5), 5 mM sodium phytate, and 50 μL enzyme preparation at 39°C for 30 min, followed by termination of reaction by adding 400 mL of an ammonium molybdate-ammonium vanadate-nitric acid mixture. After mixing and centrifugation, the absorbance was measured at a wavelength of 415 nm. One unit of phytase activity was defined as the amount of activity that liberates one micromole of phosphate per minute at 39°C.

### Dietary treatments

TG pigs and age-matched and body weight-matched non-TG pigs were used in dietary treatment experiments. The pigs were kept in individual metabolic cage. The facilities were provided with forced ventilation for thermal regulation, and each cage had one feeder and one water nipple for *ad libitum* access to feed and water. The ingredients of the selected experimental diets are listed in *Supplemental* ***table 3***. The diets were provided in pellet form.

All pigs were fed on a commercial grower diet for an adaptation period of one week in the cage, and then fed experimental diets for a pre-test period of six days prior to the start of the experiment. During the dietary treatment period, the animals were fed the experimental diets for four days. Pig feed intake was based on BW (BW × 0.04). Equal quantities of diets were added to the feeders twice daily in the morning (09:00) and afternoon (16:00), and the initial and final BW of each experimental diet were recorded.

Fecal samples were collected and processed as previously described (***Kim et al., 2005***). Feeds and dried feces were ground using a grinder, passed through a 0.425-mm size screen, and analyzed for DM (AOAC, 2005; method 930.15). Gross energy (GE) was analyzed according to ISO: 9831-1998 using a bomb calorimetry (Parr 6300; Parr Instrument Co., Louisville, KY, USA). Other nutrients in the diets and feces were analyzed using the Chinese National Standard analytical method (GB/T). The following methods were used: GB/T 6432-1994 for CP, GB/T 20806-2006 for NDF, GB/T 20805-2006 for ADF, GB/T 6434-2006 for CF, GB/T 18335-2003 for calcium, GB/T 6437-2002 for P, GB/T 6438-2007 for ash, and GB/T 21912-2008 for titanium dioxide. Apparent digestibility coefficients were calculated as described elsewhere (***Grela et al., 2011***).

### Growth performance assessment

The growth of the F2 TG pigs of both genders was compared to that of age-matched WT littermates fed on the same diets listed in ***Supplemental table 9***. A total of 23 TG boars (32.9 ± 3.80 kg) and 21 WT littermates boars (31.5 ± 2.97 kg) were grouped according to weight, and randomly allocated to seven pens fitted with MK3 FIRE feeders (**FIRE**, Osborne Industries Inc., Osborne, KS, USA). Similarly, 51 TG gilts (31.8 ± 3.55) and 31 WT gilts (30.2 ± 1.64) were randomly allocated to 11 pens that were fitted with MK3 FIRE feeders. Individual feed intake and BW were recorded when a pig with an ear transponder visited the FIRE feeders. All pigs were allowed free access to water throughout the measurement phase.

### Statistical analyses

The data were analyzed using the GLM procedure (SAS Inst. Inc., Cary, NC, USA). For the apparent total tract nutrient digestibility values, fecal nutrient output, and the growth performance, analysis of covariance (ANCOVA) was used. The BW of the tested pigs at the start of the corresponding experimental period was used as the covariable. Least square means were calculated and the differences between means were tested using Turkey-Kramer adjustment for multiple comparisons when appropriate. For salivary protein and saliva secretion by the parotid gland, one-way ANOVA followed by Duncan's multiple-comparison tests were used. For serum biochemical endpoints and saliva enzymes secretion, an unpaired two-sample *t*-test (two-tailed) was used. The level of significance was set at P < 0.05, and trends were discussed at P < 0.1.

## Acknowledgments

We thank Dr. Hao Zhang (South China Agricultural University, Guangzhou, China) for data analysis and to Dr. Stefan Moisyadi (University of Hawaii, Honolulu, HI, USA) for providing transposon plasmids. We thank Dr. Defu Zhang and Dr. Yufang Ma for providing the *cappA* gene. The National Major Project for Production of Transgenic Breeding Grant (2016ZX008006002) supported this study. We would like to thank LetPub (www.letpub.com) for providing linguistic assistance during the preparation of this manuscript.

